# A theoretical framework for proteome-scale single-molecule protein identification using multi-affinity protein binding reagents

**DOI:** 10.1101/2021.10.11.463967

**Authors:** Jarrett D. Egertson, Dan DiPasquo, Alana Killeen, Vadim Lobanov, Sujal Patel, Parag Mallick

## Abstract

The proteome is perhaps the most dynamic and valuable source of functional biological insight. Current proteomic techniques are limited in their sensitivity and throughput. A typical single experiment measures no more than 8% of the human proteome from blood or 35% from cells and tissues ^1, 2^. Here, we introduce a theoretical framework for a fundamentally different approach to proteomics that we call Protein Identification by Short-epitope Mapping (PrISM). PrISM utilizes multi-affinity reagents to target short linear epitopes with both a high affinity and low specificity. PrISM further employs a novel protein decoding algorithm that considers the stochasticity expected for single-molecule binding. In simulations, PrISM is able to identify more than 98% of proteins across the proteomes of a wide range of organisms. PrISM is robust to potential experimental confounders including false negative detection events and noise. Simulations of the approach with a chip containing 10 billion protein molecules show a dynamic range of 11.5 and 9.5 orders of magnitude for blood plasma and HeLa cells, respectively. If implemented experimentally, PrISM stands to rapidly quantify over 90% of the human proteome in a single experiment, potentially revolutionizing proteomics research.

## Main text

Despite the wealth of insights gained from increasingly routine genomic and transcriptomic studies, there remains a large gap between genomic and transcriptomic findings relationship to phenotypes. Proteomics is crucial to bridging this gap as proteins constitute the main structural and functional components of cells. However, proteome measurement technologies are significantly lagging behind next-generation DNA and RNA sequencing technologies, largely in part due to the complex nature of individual proteins and proteomes as well as the high dynamic range (∼10^9^) required for comprehensive proteomic analysis ^3^. Strikingly, roughly 10% of the proteins predicted to comprise the human proteome have never been confidently observed ^4, 5^.

Recently, single-molecule identification has been proposed as a method to analyze small samples (including single cells) and identify rare proteins ^6, 7^. Traditional bulk quantification techniques like mass spectrometry and immunoassays have been adapted towards detection of single proteins ^8, 9^. In addition, several concepts have been proposed to achieve single-molecule peptide sequencing. These all use sequential processes to determine positional information from amino acids within peptides e.g., Edman degradation ^10, 11^ or directional translocation of a peptide through a nanopore channel ^12^. However, no current method achieves both extremely high throughput and single-molecule sensitivity of intact proteins. We therefore set out to develop a fundamentally new approach to whole proteome analysis at the single molecule level.

To rapidly identify and quantify proteins at the single molecule level, we envision an experimental workflow as shown in **Fig. 1a**. Proteins are extracted from the sample and conjugated intact, but under denaturing conditions, onto a chip consisting of a hyper-dense single molecule array. On this chip, no more than one protein molecule is present per landing pad and each position is optically resolvable. Affinity reagents (antibodies, aptamers, or small proteins) are labeled with fluorophores to allow for optical detection of single-protein molecule binding events, and then passed over the chip. One affinity reagent is used per cycle, and each reagent is washed off the chip before the next one is added. Integrated fluidics and imaging allows for high resolution multi-cycle imaging of individual binding events at scale. Binding of affinity reagents to proteins produces a bind/no-bind outcome series for each protein, which can be used to infer the identity of the protein. Since there is only one protein per landing pad, direct counting can be used to quantify each protein in the sample.

**Figure 1.**
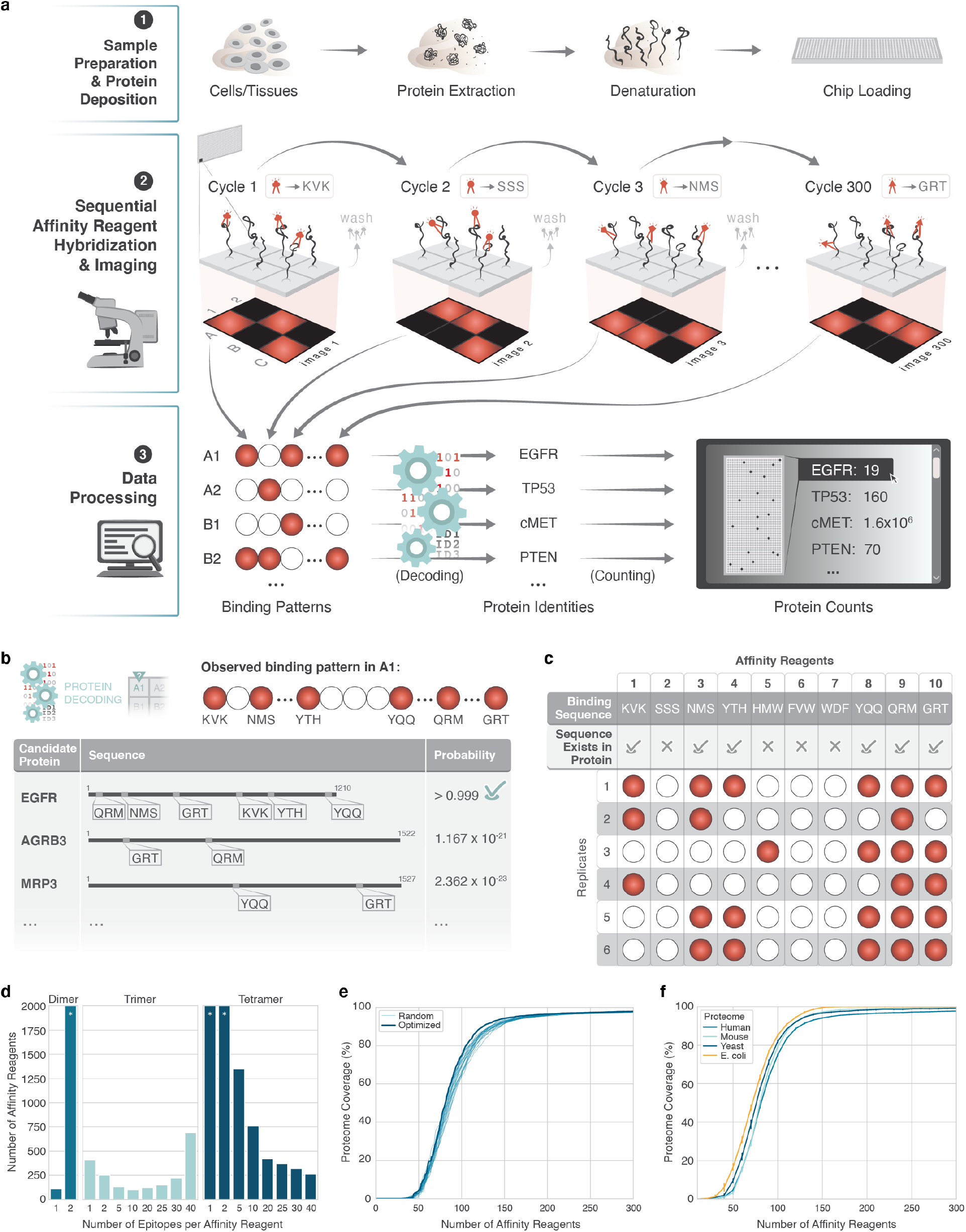
Protein Identification by Short-epitope Mapping (PrISM) potentially decodes >95% of the proteome in a wide range of organisms. **a**) The complete PrISM workflow. **b**) A depiction of protein decoding resulting in identification of the protein at location A1 as EGFR. **c**) Repeat sequential affinity reagent measurements on EGFR showing 5 unique binding patterns and one off-target binding event (rep 3, HMW). **d**) Number of affinity reagents required for 90% human proteome coverage with variation in length of epitope (dimer, trimer, tetramer) and number of epitopes bound by each multi-affinity reagent. Asterisk indicates that >2,000 affinity reagents are required. **e**) Proteome coverage achieved by measuring a panel of 300 of affinity reagents targeting trimer epitopes that were either optimized for the human proteome or randomly generated **f**) Proteome coverage for human, M. *musculus, S. cerevisiae*, and *E. coli* proteomes measured with an affinity reagent set optimized for human proteome coverage.

Identifying the tens-of-thousands of different proteins in a proteome would require a prohibitively large number of traditional highly specific affinity reagents. We therefore explored the possibility of using affinity reagents that bind short, linear epitopes (e.g., trimers) with moderate specificity, such that each affinity reagent binds many proteins. We term these reagents ‘multi-affinity’ binding reagents. While the binding of a single multi-affinity reagent is not sufficient to identify a given protein, sequential probing using a series of multi-affinity reagents can decode many proteins as we discuss below. In this approach, each new affinity reagent that is introduced in a cycle of binding and imaging provides additional evidence and gradually narrows the list of possible protein identities (**Fig. 1b**). Hereafter, we refer to our proposed approach as Protein Identification by Short-epitope Mapping (PrISM).

It would be an oversimplification to assume that there is one binding pattern or “signature” for each protein. In reality, binding is stochastic; an affinity reagent will not always bind every protein containing its target epitope when making single-molecule measurements ^13^. Furthermore, each affinity reagent may bind to off-target epitopes with similarity to the target epitope. Therefore, we anticipate measuring the same protein multiple times will likely produce a different binding pattern each time (**Fig. 1c**).

To account for this stochasticity, we developed a probabilistic binding model in which each affinity reagent binds with a primary probability to a protein containing one copy of its target epitope and with an equal or lower probability to a protein containing one copy of an off-target epitope. We began with the rather low probability of 0.5 for on-target binding to a primary epitope and 0.5 probability of binding to an off-target epitope. This is because many factors could prevent binding of an affinity reagent to its epitope, e.g., remaining protein structure due to micro-folding or incomplete denaturation, post-translational modifications and binding kinetics.

We analyzed the relationship between affinity reagent attributes (e.g. number of reagents, target epitope length, off-target binding extent and binding stochasticity) and percent of the 20,235 proteins in the SWISSPROT reference human proteome that could be identified (see Protein Sequence Databases and Protein Decoding in methods for more detail). This analysis determined that confidently identifying greater than 90% of the human proteome using affinity reagents targeting single target dimer, trimer or tetramer epitopes required 110, 410 or greater than 2,000 affinity reagent-imaging cycles (**Supplementary Table 1**). We next analyzed the impact of both target epitope lengths (dimer, trimer, or tetramer) and varied extent of off-target binding on protein identification (**Fig. 1d**). If each affinity reagent bound a single target trimer and nine additional primary biosimilar off-target trimers, 100 affinity reagents would be required to confidently identify greater than 90% of the human proteome. Each affinity reagent would bind roughly 23.7% of the total proteins; 24 binding events would be observed on average for each protein (**Supplementary Table 1**). 150 affinity reagents would be required to confidently identify greater than 97.8% of the human proteome. We found targeting tetramers would require fewer binding events to identify each protein but many more affinity reagents would be required to identify more than 90% of the proteins in the proteome. Furthermore, targeting dimers would require a similar number of affinity reagents as targeting trimers. However, any off-target binding substantially impacted the number of affinity reagent-imaging cycles required. It would likely be challenging to generate affinity reagents exclusively recognizing a single dimer. Counterintuitively, we find that less specific affinity reagents allow broader coverage of the proteome more efficiently.

As well as having primary off-target binding epitopes, affinity reagents are likely to bind to additional off-target epitopes with a lower affinity and thus a lower probability. We therefore consider an affinity reagent model (see Methods) where each affinity reagent has an additional “tail” of up to 20 secondary off-target epitopes in which binding probabilities are proportional to the sequence similarity between the target and off-target epitopes. Using this model with target epitopes selected randomly from targets present in the proteome, our simulations demonstrate that PrISM, implemented using 150 affinity reagent-imaging cycles can confidently identify 92.4% of the proteins in the human proteome. However, 300 affinity reagents are required to confidently identify ∼98% of proteins in the human proteome (**Fig. 1e**). Performance can be modestly improved when using a greedy-selection algorithm (see Methods) to select the primary target epitope for each reagent within our panel of affinity reagents (vs selecting the panel of target epitopes randomly). We used this approach to define an optimized panel of reagents in which 150 affinity reagents are able to achieve 94.2% proteome coverage and 300 affinity reagents ∼98% proteome coverage. For subsequent analyses, we used this optimized panel consisting of 300 trimer-based affinity reagents, in which each reagent bound to 1 target epitope, 9 primary off-target epitopes, and 20 secondary off-target epitopes.

To test whether PrISM could be applied across species, we used the same panel of optimized affinity reagents to simulate the analysis of proteomes from *M. musculus, S. cerevisiae*, and *E. coli* (**Fig. 1f**). Strikingly, we found minimal difference in proteome coverage between the species, indicating that, while smaller proteomes are slightly easier to decode, the primary driver of decoding performance is protein sequence diversity. Therefore, despite the stochastic nature of single molecule binding, we expect our decoding strategy is capable of identifying 98% or more of the proteins in the proteome for a wide range of organisms.

We also examined the impact of potential experimental confounders. We first considered a scenario where the probability of an affinity reagent to epitope binding is even lower than 0.5, for example due to poor binding affinity or kinetics. Even with a binding probability of 0.15, PrISM is able to identify over 85% of the human proteome using 300 affinity reagents, although this dropped to ∼55% with a binding probability of 0.1 (**Fig. 2a**). This issue could be overcome by using more affinity reagents, multiplexing affinity reagents in a single cycle (using different fluorescent labels for each probe); running affinity reagents in replicate cycles; increasing concentration and/or incubation time of affinity reagents; or attaching multiple copies of an affinity reagent to a single fluorescent particle to drive avidity. Therefore, we believe our PrISM methodology offers a viable path to massively parallel protein identification even with low affinity reagent binding probabilities.

**Figure 2.**
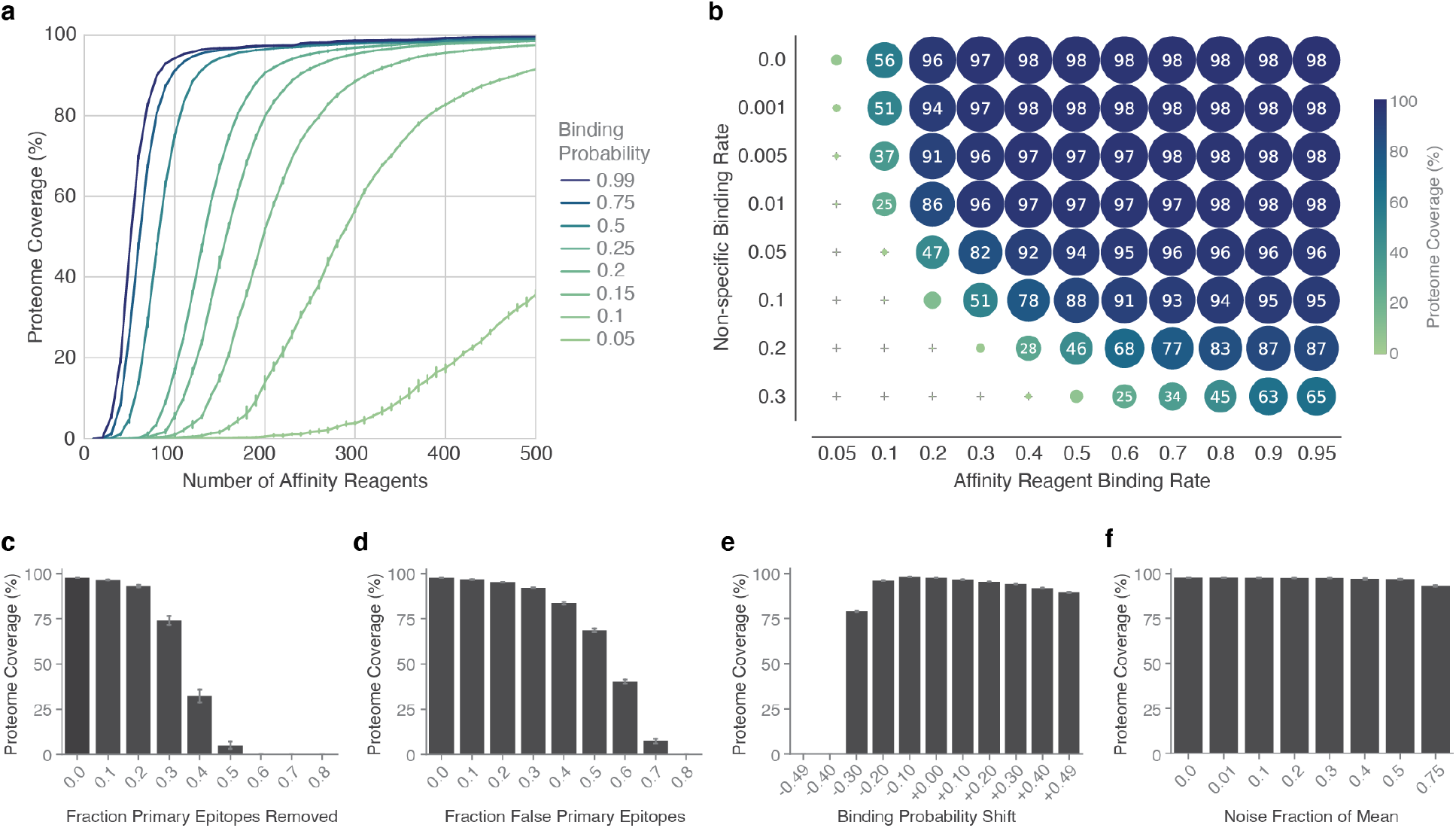
PrISM is robust to experimental confounders. **a**) Coverage of the human proteome for affinity reagents of varying binding affinity. **b**) Coverage of the human proteome for affinity reagents of varying binding affinity with non-specific binding to chip surface. Circle area is proportional to proteome coverage (also labeled on circle). **c-f**) Impact of mischaracterization of affinity reagent binding on proteome coverage for **c)** varying fraction of unknown high-affinity epitope targets, **d)** varying fraction of false high-affinity epitope targets identified, **e)** systematic measurement error in binding probability, and **f)** random measurement error in binding probability. All error bars are standard deviation across five replicates.

We next investigated the effect of non-specific binding of an affinity reagent to the chip surface close enough to a protein such that it creates a false binding signal (**Fig. 2b**). If we assume a binding probability of 0.5, then a non-specific binding rate of 0.05 or lower enables identifying >90% of the human proteome. For subsequent analyses, we assume a non-specific binding rate of 0.001. We note that the non-specific binding rate can be modulated through surface passivation, and optimization of buffers and blocking reagents.

Another source of error could be due to failures in affinity reagent characterization, i.e., the experimental determination of the target and off-target trimers and their respective binding probabilities. Such characterization can be performed in a straightforward manner using traditional epitope mapping approaches^14^. One error mode might be if an affinity reagent bound to epitopes not observed during characterization efforts leading an affinity reagent to have apparent false-positive binding to additional sites that the inference algorithm did not predict (**Fig. 2c, Suppl Fig 1a**). This impact is predicted to be small, as long as affinity reagent characterization is generally able to identify high probability (0.5) binding sites. Proteome coverage remained above 92% if up to 20% of these sites were missed. Epitopes may be falsely identified as targets during affinity reagent characterization (**Fig. 2d, Suppl Fig 1b**) leading an affinity reagent to have apparent increased false-negative binding to some epitopes. PrISM appears extremely robust to this type of error, as it achieved nearly 70% coverage even if half of all primary epitopes were incorrect. PrISM appears to be more robust to false-negatives than to false-positives in the affinity reagent model. Consequently, the techniques used to characterize affinity reagents should prioritize sensitivity over specificity. We also analyzed the impact of consistent over- or underestimation of affinity reagents’ trimer binding probabilities and find these errors have minimal impact, with the exception of large (>-0.2) underestimations of binding probability (**Fig. 2e, Suppl Fig 1c**). Performance of the PrISM analysis was also highly robust to noisy affinity reagent characterization (**Fig. 2f, Suppl Fig 1d**), indicating that affinity reagent characterization need not be perfect, and that our method will tolerate variability in affinity reagents to protein binding that may arise from other experimental confounders. In summary, we are confident the PrISM methodology is robust to errors in affinity reagent characterization.

Complex mixtures, such as cell lysates and blood plasma samples are consistently challenging for current proteomics methods; lysate and plasma protein abundances span a range of more than 9-12 orders of magnitude ^15^ and typical mass spectrometry-based approaches identify less than 8% of the blood plasma proteome^1^ and 35% of the cellular proteome. To evaluate the theoretical performance of PrISM on lysate and plasma, we ran simulations assaying cell lysates and un-depleted blood plasma sample with 300 affinity reagents on model chips with 10^10^ landing pads. The simulation modeled running the same sample with 5 technical replicates. Random noise in affinity reagent to trimer binding probability simulated variability in affinity reagent binding across technical replicates.

On average, simulations executing PrISM with a 10^10^ landing pad chip on a HeLa sample demonstrated deposition of at least 1 copy of 97.8% of proteins present in the HeLa sample leading to detection of 92.6% of the proteins in the human proteome. The detected proteins spanned 9.5 orders of magnitude dynamic range (**Fig 3a, Suppl Fig 2**). With the same simulation run on a plasma sample, the dynamic range of detected proteins is >11.5 orders of magnitude (**Fig 3b**). However, due to the high sample dynamic range of plasma this wide deposition dynamic range represents a smaller fraction (66%) of the proteome than the HeLa sample. Almost all proteins were quantified with high specificity (**Supp Fig 3**); more than 99.9% of the measured proteins had quantitative specificity >90% (i.e., >90% of identifications of the protein were true positives). Increasing the number of landing pads from 10^10^ to 10^11^ or 10^12^ does not notably impact the deposition of a HeLa sample (**Suppl Fig 4a**). However, it would increase plasma proteome deposition on the chip from 66% to 79% or 92% respectively (**Suppl Fig 4b**). Experimentally, sample dynamic range could be further compressed by depleting the most abundant proteins in a plasma sample, e.g., using an affinity column. A plasma sample modeled with 99% depletion of the top 20 proteins had 73%, 85%, and 97% proteome deposition on a chip of 10^10^,10^11^, and 10^12^ landing pads, respectively (**Suppl Fig 4c**). In both cell lysate and plasma samples, we confidently detected >90% of proteins deposited on a 10^10^ chip, suggesting the primary limitation of detection dynamic range is not the ability to decode proteins, but is rather the ability to deposit low concentration proteins onto the chip. These analyses suggest that PrISM has the potential to quantify the vast majority of the human proteome across a wide dynamic range within a single experiment.

**Figure 3.**
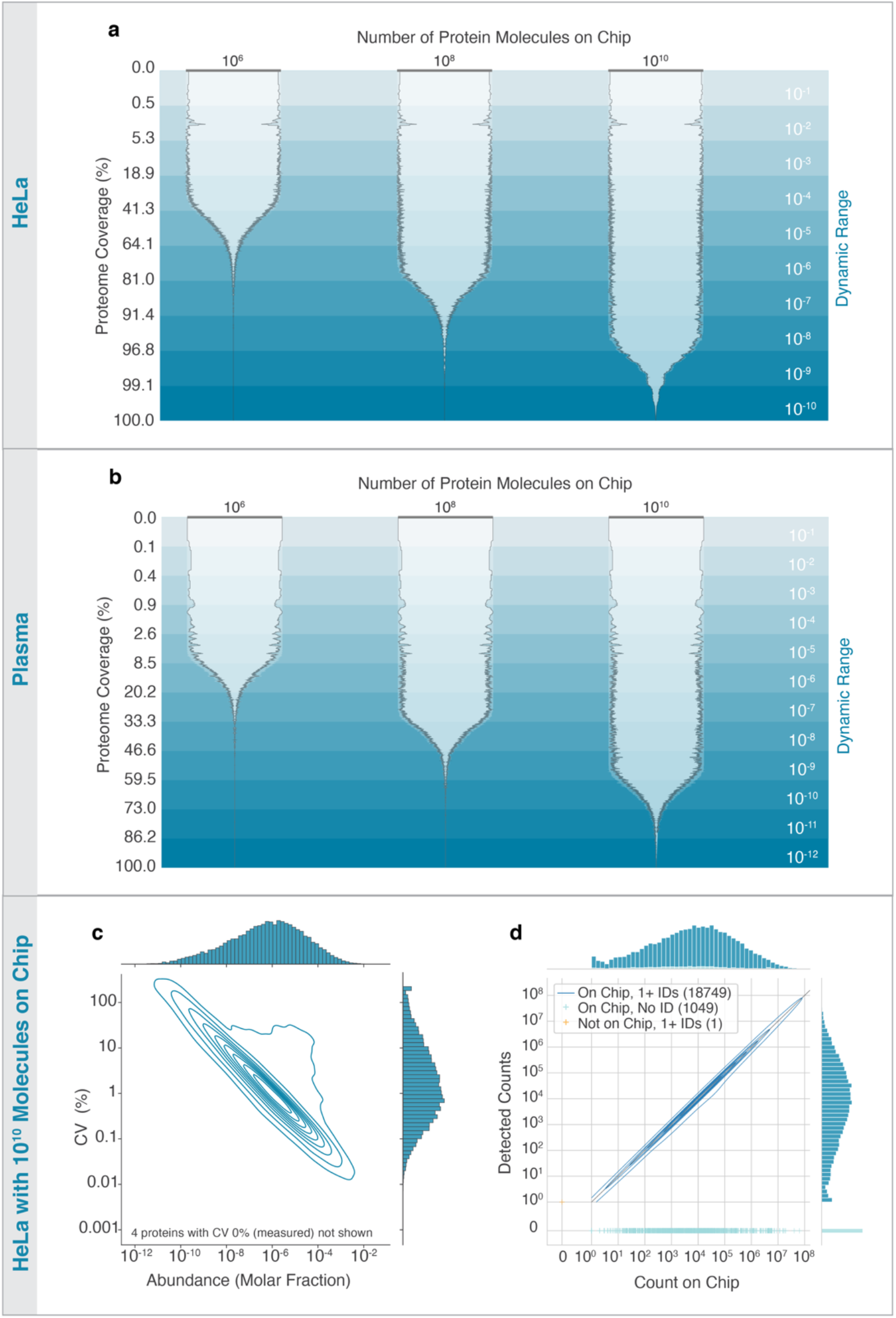
Simulated proteome experiments for HeLa cells and plasma demonstrate large dynamic range of detection and high coverage. **a)** Dynamic range of protein quantification for HeLa cell lysate with varying chip size. Data are plotted in order of decreasing protein abundance from top to bottom. Dynamic range is the protein abundance divided by the most abundant protein in sample. The outer width of the contours indicates the percentage of proteins at that abundance deposited on the chip (one or more copies). The inner width of the contours indicates the percentage of proteins at that abundance detected by PrISM. Percentages are computed over a rolling window of 51 proteins. Horizontal gray bars indicate 100%. Panels c,d were computed on a 10^10^ chip. **b)** Dynamic range of protein quantification for an undepleted plasma sample, with varying chip size. **c)** Reproducibility of quantification (coefficient of variation computed across five replicates) compared to protein abundance for HeLa cell lysate as contour plots (density iso-proportional contours) with marginal histograms. **d)** Concordance of quantity of proteins (number of copies identified) measured by PrISM with true count of protein on chip for a single experimental replicate of HeLa cell lysate as contour plots (density iso-proportional contours) with marginal histograms.

We assessed measurement reproducibility across the 5 technical replicates in HeLa samples (**Fig. 3c**) and plasma samples (**Suppl. Fig 5a, 5b**). 72% and 85% of proteins within the top 5 orders of magnitude had a CV <1% in HeLa and plasma, respectively. As modeled, the contributors to irreproducibility were stochastic variation in affinity reagent binding and protein deposition as well as variation in affinity reagent binding characteristics. While these estimates do not consider many factors of experimental variability such as sample preparation and biological variability, they demonstrate the potential of the analytic platform and decoding algorithm to contribute minimal variation relative to more common sources of variation (**Suppl Fig 6a, 6b**).

Detected protein counts correlated with the number of proteins on the chip (**Fig. 3d, Suppl Fig 5b**). 85% of cell lysate proteins and plasma proteins had a fold change error in detected counts relative to counts on the chip within +/-10% (**Suppl Fig 7**) indicating that protein sensitivity is broadly uniform. In some cases, proteins with only a single copy on a chip were detected. Some proteins were substantially under-counted due to sequence similarity to other proteins in the sequence database. The linear nature of detection count vs. counts on chip indicates that dynamic range can be extended further by increasing spots on the chip (10^11^ or higher) or by running a sample on multiple chips.

In conclusion, PrISM is a single-molecule species agnostic protein identification approach that enables analysis of nearly the entire human proteome in a single experiment. PrISM has important advantages over other proteome analysis methods. Amongst emerging single-molecule peptide sequencing approaches, it is unique in taking a non-destructive affinity reagent approach, rather than a chemically intensive or cleavage-based sequencing approach. PrISM is robust to both false negatives (i.e., failure of an affinity reagent to bind its epitope) and false positives (i.e. affinity reagents binding to unexpected epitopes) and has been broadly optimized for promiscuous affinity reagents. Therefore, PrISM turns a perceived weakness of affinity-based proteomic approaches into a strength. PrISM is scalable to full proteome quantification over wide dynamic ranges. The uniform protein sensitivity of PrISM simplifies protein quantification to direct counting of single-molecule identifications. This approach stands in strong contrast to quantification in bottom-up mass spectrometry approaches, which must address factors like variable peptide ionization efficiency and the assembly of peptide measurements into quantitative information on proteins. Furthermore, by using intact proteins, PrISM avoids the loss of information (such as proteoforms) that limits peptide-centric approaches. Additionally, focusing on intact proteins partially mitigates the dynamic range challenge, as complex mixtures are not made more complex through digestion of each protein into 10s-100s of peptides. PrISM may be readily extended to the study of post-translational modifications through the use of affinity reagents targeting altered residues and motifs (e.g. phospho-specific affinity reagents). Furthermore, as an intact protein method, PrISM may be readily extended to analysis of proteoforms. PrISM has some limitations: *de novo* protein identification is currently not possible, as our approach relies on a sequence database. Therefore, applications to studying the immunopeptidome and gut meta-proteome are likely best suited to other approaches. Additionally, distinguishing highly similar proteins (including those differing by mutation) will be challenging. However, this may be mitigated by developing panels of affinity reagents targeting conserved or mutated regions in proteins of interest.

Future work required to implement PrISM experimentally will include developing and characterizing appropriate affinity reagents and setting up a sensitive and rapid imaging platform. As the dynamic range of the method is directly related to the number of intact protein molecules measured, imaging and cycle speed will be critical. With 300 affinity reagents, introduced two at a time, and measured with two-color imaging, an affinity reagent-imaging cycle time of approximately 10 minutes would make it possible to profile ten-billion protein molecules within about a day. It may be further possible to increase sample throughput by titrating number of molecules measured, or with additional multiplexing.

A successful experimental implementation of PrISM will provide a user-friendly, rapid, ultrasensitive, and reproducible method to analyze and quantify proteomes, even from single cells. It would open a path to countless new opportunities in scientific discovery, not only in basic research but also in clinical research, including molecular diagnostics and biomarker discovery.

## Methods

### Protein Sequence Databases

Protein sequence databases were downloaded from UniProt (www.uniprot.org). For each species, the “reference” proteome was selected by including “reference:yes” in the search query string for proteomes. The reference proteome was then filtered to only include Reviewed (Swiss-prot) sequences (query string “reviewed:yes”). The sequence data were then downloaded in uncompressed .fasta format (canonical sequences only). Specific proteomes and filter strings used were:

- *E. coli* (strain K12): reviewed:yes AND organism:”Escherichia coli (strain K12) [83333]” AND proteome:up000000625 (downloaded 6/30/2021)
- *S. cerevisiae* (s288c): reviewed:yes AND organism:”Saccharomyces cerevisiae (strain ATCC 204508 / S288c) (Baker’s yeast) [559292]” AND proteome:up000002311 (downloaded 6/30/2021)
- *M. musculus* (c57bl): reviewed:yes AND organism:”Mus musculus (Mouse) [10090]” AND proteome:up000000589 (downloaded 6/30/2021)
- *H. sapiens*: reviewed:yes AND organism:”Homo sapiens (Human) [9606]” AND proteome:up000005640 (downloaded 7/6/2021)

The proteomes were further processed to remove any duplicated sequences and any sequences not entirely composed of the 20 canonical amino acids. Further, sequences of length 30 or less were removed from each FASTA.

### Modeling Affinity Reagent to Protein Binding

An affinity reagent targeting epitopes of length *k* (e.g., for a trimer, *k* = 3) was modeled by assigning a binding probability *θ* to each unique target epitope *j* of length *k* recognized by the reagent. Further, a protein non-specific binding rate was assigned *p*_*nsbepitope*_ representing the probability of the affinity reagent binding to any epitope in a protein non-specifically. Given the primary sequence for a protein of length *M*, the probability of an affinity reagent binding to the protein was computed as follows:

First compute the probability of a specific binding event happening:

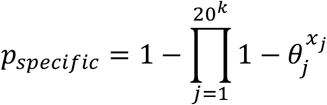

**with:**

- ***X*: the count of each epitope *j* in the protein sequence**
  - *X* = {*x*_1_, *x*_2_, *x*_3_…} with *x*_*j*_ ∈ 𝕫^∗^

- ***θ*: the binding model parameters. A vector of probabilities of the affinity reagent binding to each recognized epitope**
  - *θ* = {*θ*_1_ *θ*_2_, *θ*_3_, …} with 0 ≤ *θ P*_*nsbepitope*_ ≤ 1

Next, compute the probability of a non-specific protein binding event happening:

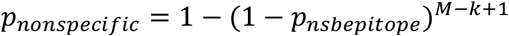

**with:**

- ***P***_***nsbepitope***_**: the probability of the affinity reagent non-specifically binding to any epitope in the protein**
  - 0 ≤ *P*_*nsbepitope*_ ≤ 1
- *M*: the length of the protein sequence
- *k*: the length of the linear epitope(s) recognized by the affinity reagent

The probability of the affinity reagent binding to the protein and generating a detectable “light-up” event is the probability of 1 or more specific or non-specific binding events occurring:

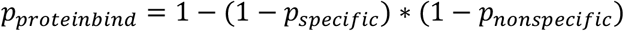

When noted, the probability of binding to each protein was adjusted to account for additional random “surface” non-specific binding. That is, binding of an affinity reagent to the chip close enough to the location of the protein to generate a false-positive binding event. The prevalence of surface NSB is defined as a probability 0 ≤ *p*_*surfacensb*_ < 1 of such a surface NSB event occurring during the acquisition of a single affinity reagent measurement at a single protein location on the chip. The adjusted probability of a protein binding event taking into account surface NSB is:

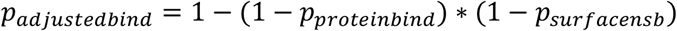

### Biosimilar Affinity Reagent Model

Unless specifically noted, affinity reagents were modeled using a “biosimilar” model. In this model, an affinity reagent targets a specific epitope which it binds with probability 0.5. The affinity reagent also binds nine additional primary “off-target” epitopes with probability 0.5 that are “biosimilar” to the targeted epitope. Biosimilar targets were selected by computing a pairwise similarity score of the target epitope to every other possible epitope of the same length. The similarity score was computed by summing up the BLOSUM62 similarity between the pair of residues at each sequence location. For example, if computing the similarity of a trimer SLL with trimer YLH, the score would be BLOSUM62(S,Y), + BLOSUM62(L,L) + BLOSUM62(L,H). With all pairwise similarity scores computed, the top nine most similar epitopes to the target were selected as the primary “off-target” epitopes. In the case of a tie where multiple potential off-target epitopes had the same score, a random epitope was selected. In addition to the target epitope and nine off-target epitopes, up to 20 additional secondary biosimilar off-target sites of lower binding probability were added to the affinity reagent. The 20 secondary off-target sites bind to the next 20 most biosimilar epitopes beyond the ones already included in the affinity reagent model. These 20 additional sites have a probability computed as

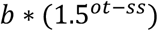

with:

- *b* = binding probability of the affinity reagent to its target
- *ot* = BLOSUM62 similarity score between affinity reagent target and this off-target site
- *ss* = BLOSUM62 similarity score between affinity reagent target and itself

If any of these additional off-target sites had binding probability that is less than the affinity reagent epitope non-specific binding rate, it was not included. The epitope non-specific binding probability was set at 2.45*x*10^−8^

### Simulation of Stochastic Affinity Reagent Binding

To simulate binding of a series of affinity reagents to a single protein, the binding probability *θ*_*i*_ of each affinity reagent *i* to the protein was first determined using the methods described in <u>Modeling Affinity Reagent to Protein Binding</u>. To simulate the outcome of the binding for each affinity reagent, a single random draw was taken from the Bernoulli distribution parameterized by *θ*_*i*_. An outcome of 1 is binding, an outcome of 0 is no binding.

### Protein Decoding

#### Overview

The protein decoding algorithm analyzes a series of affinity reagent binding measurements acquired on an unknown protein and determines the most likely identity of that protein among a set of candidates. The most likely protein identity is the one most compatible with the observed binding measurements. This compatibility is determined based on binding models for each affinity reagent in the experiment, which are used to estimate how likely each affinity reagent is to bind each potential protein. A strong candidate protein is one where most of the observed binding events are consistent with affinity reagents likely to bind that protein. A weak candidate protein will have many instances where binding is observed for affinity reagents that are not expected to bind the candidate. The strongest candidate protein is deemed the most likely identity for the unknown protein and confidence in this identification is computed as a relative measure of the compatibility of the most likely protein compared to all the other candidates.

#### Inputs

**The inputs to the decoding algorithm are:**

- **Binding data:** *D* = [*d*_1_ *d*_2_, *d*_1_… *d*_*N*_] with *d* ∈ {0(*nobind*),1(*bind*)}. A sequence of binding measurements, one for each affinity reagent to an unknown protein.
- A **sequence database** of length *M* containing the primary sequence and name of each potential protein that may be present in the sample (e.g., the human protein sequence database described in <u>Protein Sequence Databases</u>)
- A parameterized **binding model** for each of the *N* affinity reagents used in the experiment (see <u>Modeling Affinity Reagent to Protein Binding</u>) where each affinity reagent binds to epitopes of length *k*. For a trimer model, *k* = 3 and there are 20^k^ (8000) possible unique epitopes the reagent could bind to.
- An (optional) **surface non-specific binding rate** (*r*) describing the probability of a surface non-specific binding event happening at any one landing pad in any given cycle.

### Binding Probability Calculations

An *M* × *N* binding probability matrix **B** was computed describing the probability of each affinity reagent binding to every possible candidate protein with an entry in the matrix *b*_*i,j*_ being the probability of affinity reagent *j* binding to candidate protein *i*. These probabilites were computed using the methods described in <u>Modeling Affinity Reagent to Protein Binding</u>.

Next, the *M* × *N* matrix **U** with adjusted non-binding probabitities for each affinity reagent to each protein was computed as follows:

- Compute *S* = [*s*_1_ *s*_2_, *s*_1_, … *s*_3_] where *s*_*i*_ = *protein*_*i*_ *length* − *k* + 1
- Compute *F* = [*f*_1_ *f*_2_, *f*_1_, … *f*_20_^*k*^] the relative frequency of every possible unique epitope among the set of all candidate protein sequences where 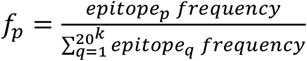
- Compute *A* = [*a*_1_ *a*_2_, *a*_1_, … *a*_*N*_] the vector of average epitope non-binding probabilities for the affinity reagents. A value *a*_*j*_ in *A* is the probability of the affinity reagent not binding to an epitope, averaged over all 20^*k*^ epitopes and weighted by the relative frequency of each epitope in the candidate protein database. 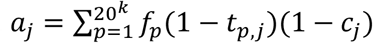 where *t*_*p,j*_ is the probability of affinity reagent *j* binding to epitope *p* and *c*_*j*_ is the probability of a non-specific protein binding event happening for affinity reagent *j*.
- Compute **U** where 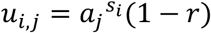 is the adjusted probability of affinity reagent *j* not binding to protein *i* (*r* is the surface NSB rate)

Adjusted non-binding probabilities were computed in this manner (as opposed to **U** = 1 − **B**) to avoid any single non-binding event having an outsized impact on a protein. Our rationale was that there are numerous difficult-to-predict reasons why an affinity reagent may not bind to a specific site (e.g., protein structure, PTMs) and so the total number of non-binding events should be considered more than the specific identity of the observed non-binding events.

### Decoding

A vector of likelihoods for each protein in the candidate database was computed by multiplying the likelihoods of each observed binding event.

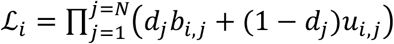

The protein of highest likelihood was selected (if there was a tie for top protein, one of the top proteins was selected randomly):

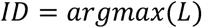

The probability of the ID being correct is the likelihood of the top protein divided by the sum of the likelihood of all other proteins:

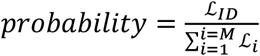

The protein *ID* and *probability* are the outputs for the decoding process performed on a single unknown protein.

### Calculation of Theoretical Proteome Coverage

To compute proteome coverage requires definition of a set of affinity reagents defined as in <u>Modeling Affinity Reagent to Protein Binding</u>. Binding of the affinity reagents was simulated for each protein (see <u>Simulation of Stochastic Affinity Reagent Binding</u>) in the human proteome as defined in section <u>Protein Sequence Databases</u>. The binding data were then passed to the decoding algorithm along with the definition of the affinity reagents, and the FASTA sequence database. The output of the decoding algorithm is a single protein identification for each simulated protein and an estimated probability of that identification being correct. To compute the fractional coverage, the number of proteins identified above a true false discovery rate (FDR) threshold of 1% (see <u>Computing and Thresholding on False Discovery Rate</u>) was divided by the total number of proteins simulated. The percent coverage was computed by multiplying fractional coverage by 100. This method was applied for all analyses except for modeling of cell, plasma, and depleted plasma samples which used the method described in <u>Quantitative Statistics</u>.

### Computing and Thresholding on False Discovery Rate

Given a list of decode protein identities (protein identity and associated probability), the FDR was computed by first annotating each protein identification as correct or incorrect based on its match to the true identity of that protein in the simulation. For each unique identification probability in the list, the FDR was computed as the fraction of proteins at that probability or lower that were incorrectly identified. To threshold on FDR, the lowest probability score threshold with FDR less than the desired FDR was determined. Identifications at this probability score or higher satisfied the FDR criterion and were considered “identified” at the desired FDR threshold.

### Demonstration of Stochastic Binding

Stochastic binding of a sequence of ten affinity reagents to protein EGFR was simulated six times. Affinity reagents with binding sequence present in EGFR have a 0.5 probability of binding and those without a binding sequence in EGFR have 0.05 probability of binding. Binding was simulated as described in <u>Simulation of Stochastic Affinity Reagent Binding</u>.

### Evaluation of Affinity Reagent Requirements for Efficient Decoding

Affinity reagents with various target epitope lengths (2, 3, or 4 i.e., dimer, trimer, tetramer) with varying numbers of primary off-target sites were modeled. In each case, the target binding probability was 0.5. “Number of Epitopes per Affinity Reagent”=1 represents affinity reagents targeting a single epitope, with no primary off-target sites. Other scenarios were modeled with the affinity reagents having some number of primary biosimilar (see <u>Biosimilar Affinity Reagent Model</u>) off-target sites. For example, an affinity reagent labeled as targeting “5” epitopes has binding affinity for its target and four primary off-target sites. Affinity reagents did not have any secondary off-target sites (<u>Biosimilar Affinity Reagent Model</u>). The targets of affinity reagents were selected randomly from targets present in the proteome. There was no requirement for off-target binding sites being present in the proteome.

To determine the number of affinity reagents required to achieve 90% coverage of the proteome, binding of an excess of affinity reagents (i.e., more than required for 90% coverage) was simulated to each protein in the proteome. For any number of affinity reagents N, the proteome coverage may be computed using the first N affinity reagents in the set. The number of affinity reagents required to achieve 90% proteome coverage is the lowest N with coverage at or exceeding 90%. The values of N tested were in increments of 10.

With the number of affinity reagents (N) required for 90% coverage computed, the number of binding events observed for each simulated protein was recorded, and the mean of these values reported as the “Average Number of Binding Events per Protein”. Additionally, the percent of proteins generating a binding event for each affinity reagent was recorded, and the mean of these values is reported as the “Percent of Proteins Bound Per Affinity Reagent”.

### Selection and Evaluation of Optimal Affinity Reagent Trimer Targets

The standard biosimilar affinity reagent model (see <u>Biosimilar Affinity Reagent Model</u>) was used in this analysis with trimer-targeting affinity reagents. One set of “optimal” affinity reagent targets was computed by using a greedy-selection algorithm to estimate the optimal set of 300 targets to achieve high proteome coverage with as few affinity reagents as possible. Additionally, 20 sets of 300 targets were selected randomly among trimers present in the proteome (excluding any trimers containing a cysteine). Proteome coverage for each of the 21 affinity reagent sets was evaluated as described in <u>Calculation of Theoretical Proteome Coverage</u>. Proteome coverage was also evaluated for multiple first-N reagent subsets of each affinity reagent set to evaluate scaling of proteome coverage with number of affinity reagents used.

The optimal set of trimer targets is chosen as described below:

1. Initialize an empty list of selected affinity reagents (AR).
2. Initialize a set of candidate ARs (e.g., a collection of 6,859 ARs, each targeting a unique trimer without a cysteine in it).
3. Select a set of protein sequences to optimize against. In this case, all human proteins in the UniProt reference proteome are used.
4. Repeat the following until the desired number of ARs has been selected:
  a. For each candidate AR:
    i. Simulate binding of the candidate AR against the protein set.
    ii. Perform decoding for each protein using the simulated binding measurements from the candidate AR and the simulated binding measurements from all previously selected ARs.
    iii. Calculate a score for the candidate AR by summing up the probability of the correct protein identification for each protein determined by protein inference.
  b. Add the AR with the highest score to the set of selected ARs, and remove it from the candidate AR list

### Evaluation of Proteome Coverage in Multiple Organisms

Proteome coverage was assessed for four different organisms using the 300 affinity reagents targeting the optimal trimer set (<u>Selection and Evaluation of Optimal Affinity Reagent Trimer Targets</u>) designed against the human proteome. Sequence databases for each organism are described in <u>Protein Sequence Databases</u>. For each organism, binding was simulated using an affinity reagent epitope binding affinity of 0.5 for each affinity reagent against each protein in the sequence database for that organism. The binding data were then decoded using the appropriate sequence database for the organism and proteome coverage assessed as described in <u>Calculation of Theoretical Proteome Coverage</u> using various first-N subsets of the 300 affinity reagent set. For example, to compute coverage at 100 affinity reagents for a given organism, only data from the first 100 of the 300 affinity reagents total were considered when decoding.

### Application of Noise to Affinity Reagent Binding Probabilities

A method was devised to model random perturbations in affinity reagent binding characteristics. The method applies random “noise” to the trimer (or other short linear epitope) binding probabilities while maintaining probabilities bound between 0 and 1. Given a binding probability *p* a perturbed probability was determined by drawing a sample from the distribution:

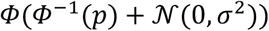

where:

- *𝒩* is the normal distribution
- *σ*^2^ is a parameter used to tune the severity of the perturbation
- *Φ* is the cumulative distribution function of the standard normal distribution

The parameter *σ*^2^ was set such that the mean absolute deviation (MAD) of the distribution divided by the trimer probability *p* is equal to a desired target. This tuning parameter will be referred to as the “fractional MAD”. The fractional MAD was used to tune the noise due to its conceptual similarity to the coefficient of variation (standard deviation divided by mean) often used to describe measurement noise or reproducibility for normally-distributed measurements.

A numerical approximation approach was used to find the value of *σ*^2^ for a probability *p* that results in the desired fractional MAD. First, given *p* and the desired fractional MAD, the target *MAD* was computed as *fractional MAD* ∗ *p*. A function *optim* was defined which, given *p*, the target *MAD*, and a proposed *σ*^2^ value generates 10,000 random samples from the noise distribution parameterized by *p* and *σ*^2^ and returns the absolute value of the difference between the *MAD* of the 10,000 random samples and the target *MAD*. The *minimize_scalar* function from the *scipy* Python package was used to estimate the value of *σ*^2^ which minimizes this function. This process was repeated 50 times, and the median optimal *σ*^2^ among the 50 trials is taken as the appropriate value to generate a noise distribution with the desired *MAD*.

### Modeling of Experimental Confounders

#### Poor Binding Affinity

Proteome coverage was assessed using the 300 affinity reagents targeting the optimal trimer set (<u>Selection and Evaluation of Optimal Affinity Reagent Trimer Targets</u>) binding to each unique protein in the human proteome. However, the affinity reagents were modeled with a variety of target epitope binding rates ranging from 0.01 to 0.99 to simulate varying affinity reagent binding affinity. Proteome coverage was assessed as described in <u>Calculation of Theoretical Proteome Coverage</u> using various first-N subsets of the 300 affinity reagent set to model the relation between number of affinity reagents used and proteome coverage. Binding simulation and decoding were repeated five times to generate replicate analyses.

#### Non-Specific Binding to Chip Surface

Proteome coverage was assessed with varying combinations of affinity reagent binding affinity and non-specific binding rate. In every case, 300 affinity reagents targeting the optimal trimer set (<u>Selection and Evaluation of Optimal Affinity Reagent Trimer Targets</u>) were used. However, the affinity reagents were modeled with a variety of target epitope binding rates ranging from 0.05 to 0.95 to simulate varying affinity reagent binding affinity and also varying “surface” non-specific binding rates ranging from 0 to 0.3. After modeling binding with surface NSB, proteome coverage was computed as described in <u>Calculation of Theoretical Proteome Coverage</u>.

#### Missed Trimers During Affinity Reagent Characterization

Binding measurements for each of the set of optimal affinity reagents (see <u>Simulation of Stochastic Affinity Reagent Binding</u>) were generated against each of the proteins in the human FASTA database (<u>Protein Sequence Databases</u>) with a surface NSB rate of 0.1% (<u>Non-Specific Binding to Chip Surface</u>). Prior to decoding the binding measurements to generate protein IDs, the affinity reagent models were “corrupted” by removing a fraction of primary epitopes. Such a corruption could occur in an experimental setting if the approach used to determine the epitopes that an affinity reagent binds to “missed” some number of epitopes. The corrupted affinity reagent models were used when decoding the binding measurements to generate protein IDs and were expected to reduce decoding performance. The severity of the corruption was modulated by adjusting the percentage of primary epitopes that were missed. To model 20% of primary epitopes being missed, a random 20% of the primary epitopes (among all affinity reagents collectively) were selected for removal. Because the optimal affinity reagents have ten primary epitopes, this means that on average, two primary epitope were missed in each affinity reagent, although some may have more than one removed and others may have none removed due to random chance. In some analyses, a percentage of secondary epitopes were also removed in a similar manner.

#### False Identification of Trimer Sites During Affinity Reagent Characterization

Similar to <u>Missed Trimers During Affinity Reagent Characterization</u>, binding of affinity reagents to proteins in the proteome was simulated with surface NSB 0.1% and affinity reagent models were corrupted prior to decoding. For this analysis, “false positive” epitopes were added to the affinity reagents prior to decoding. This simulates a scenario where the approach used to characterize the epitopes bound by each affinity reagent falsely identifies some number of trimer epitopes which the affinity reagent does not bind to. The severity of the corruption was modulated by adding false primary epitopes such that the complete set contained a specific percentage of false sites. For example, 20% false sites means that false primary epitopes were added until 20% of the primary epitopes among the affinity reagent set were false. The extra epitopes were randomly distributed among the affinity reagents. The trimer identities of the extra epitopes were selected randomly with replacement. In some analyses, secondary epitopes were also impacted by this corruption. Any added secondary epitopes must not match an existing or added primary epitope. For example, an affinity reagent targeting the primary epitopes HNW, HDW, and HHW and secondary epitopes HRW, and HGW could have LWW added as either a corrupting primary or secondary epitope, but HGW could only be added as a corrupting primary epitope, in which case its binding probability would be updated to that of a primary epitope.

#### Consistent Over- or Under-Estimation of Affinity Reagent Trimer Binding

Similar to <u>Missed Trimers During Affinity Reagent Characterization</u>, binding of affinity reagents to proteins in the proteome was simulated with surface NSB 0.1% and affinity reagent models were corrupted prior to decoding. In this analysis, epitope binding probabilities were adjusted to be systematically higher or lower than the true values. This models a situation where the affinity reagent characterization approach determines the correct trimer epitopes targeted by the affinity reagent, but systematically over or under-estimates the strength of binding (modeled by binding probability). The manipulation entails applying some fold-change shift to the binding probability of the epitopes such that the primary epitopes of the affinity reagent are shifted by a desired amount. For example, to model a shift of +0.25 for an affinity reagent with true primary epitope binding probability of 0.25, the binding probability of every epitope of the affinity reagent is multiplied by 2. In this case a primary epitope with true binding probability of 0.25 will be assumed to bind with a probability of 0.5 when performing decoding. Similarly, this same multiplicative shift may be applied to secondary binding epitopes. For example, a secondary epitope with binding probability 0.2 would then have binding probability 0.4. Similarly, adjustments may be made which adjust binding probabilities to be less. In some analyses, the severity of the corruption was modulated by only corrupting a fraction of the affinity reagents. For example, 50% of the affinity reagents may be impacted meaning half of the affinity reagents have a systematic error in their binding probabilities while the rest are not impacted.

#### Noisy Affinity Reagent Characterization

Similar to <u>Missed Trimers During Affinity Reagent Characterization</u>, binding of affinity reagents to proteins in the proteome was simulated with surface NSB 0.1% and affinity reagent models were corrupted prior to decoding. In this analysis, random “noise” was applied to the characterized epitope binding probabilities. The random noise was applied to a random fraction of the affinity reagents in the set. For any affinity reagent impacted by noise, all primary and secondary epitopes were subjected to some degree of noise as well as the affinity reagent non-specific binding rate. The binding probabilities were perturbed according to the method described in <u>Application of Noise to Affinity Reagent Binding Probabilities</u> with the amount of noise ranging between *fractional MAD* 0 and 0.75.

### Simulation of Cell-Line and Plasma Experiments

#### Protein Abundance Database Processing

The protein composition of each sample was modeled using protein abundances downloaded from PaxDb v4.1 ^16^. Specifically, plasma protein abundances were from the “H.sapiens - Plasma (Integrated)” dataset (https://pax-db.org/downloads/4.1/datasets/9606/9606-PLASMA-integrated.txt downloaded September, 2021). Cell-line abundances were from the dataset “H.sapiens - Cell line, Hela, SC (Nagaraj, MSB,2011)” (https://pax-db.org/downloads/4.1/datasets/9606/9606-hela_Nagaraj_2011.txt built from high resolution mass spectrometry analysis of HeLa cells^17^. The identities of proteins in the PaxDb data were mapped to the identities of proteins in the UniProt human protein sequence database (see <u>Protein Sequence Databases</u>) using the PaxDb to UniProt mapping available from the PaxDb maintainers available at https://pax-db.org/downloads/4.1/mapping_files/uniprot_mappings/full_uniprot_2_paxdb.04.2015.tsv.zip (downloaded September, 2021). Any proteins present in the PaxDb database that could not be mapped to the UniProt sequence database were removed from the sample. 4,342 of 4,492 entries (97%) in the plasma database were successfully mapped with no unmapped protein comprising more than 1% of the sample. 8,554 of 8,817 entries (97%) in the cell database were successfully mapped with no unmapped protein comprising more than 1% of the sample. In some cases, more than one entry in a PaxDb database mapped to a single UniProt identifier in the sequence database. In these cases, only the first entry was retained. In the plasma database, 99 database entries were dropped as a result of this operation (4,243 entries remained). In the cell-line database, 145 entries were dropped (8,409 entries remained). Neither of these operations dropped any entries comprising more than 1% of the corresponding sample. 25, and 97 proteins with abundance 0 were removed from the plasma and cell-line database, respectively. After filtering, the abundance databases were normalized to sum to 1.

#### Imputation of Protein Abundances (Plasma)

Abundances were imputed for proteins in the human protein sequence database not represented in the modeled plasma sample (<u>Protein Abundance Database Processing</u>). This process resulted in a “complete” plasma sample containing 20,235 proteins with 12 orders of magnitude in dynamic range of abundance. The distribution of abundances in the complete plasma sample was modeled as a semi-Gaussian distribution ^18^:

Let *f*(*x*|*μ, σ*) be the normal distribution probability density function with mean *μ* and standard deviation *σ* evaluated at *x*

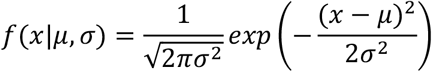

Let:

- *A*_*max*_ be the highest protein abundance in the modeled plasma sample pre-imputation
- *σ*_*p*_ = 1.2
- *μ*_*p*_ = *log*_1_(*A*_*max*_) − 5*σ*_*p*_
- 𝓁 = *log*_1_(*A*_*max*_) − 12

Let *g*(*a*) be a function proportional to the probability density of the semi-Gaussian distribution at abundance *a*.

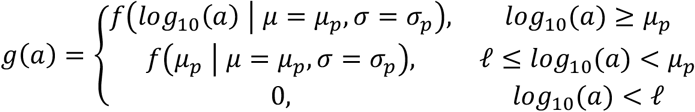

Next, a probability density function for abundances of proteins needing to be imputed was estimated. We set a threshold for “high-abundance” proteins *t* = *A*_*max*_ − 4 and reasoned that any protein with *log*_1_(*abundance*) > *t* present in the “complete” plasma sample would be accurately represented in PaxDb (i.e., not impacted by detection bias). The probability density of the PaxDb proteins was estimated by computing a histogram (50 bins) on their log-10 transformed abundances and normalizing the values at each bin such that the total area of the histogram is 1.

A scaling factor *α* was computed to adjust the high-abundance tail of the complete sample abundance distribution *g*(*x*) to match the probability density of protein abundances > *t* in PaxDb:

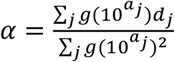

**with:**

- {*a*_0_, *a*_1_ *a*_2_, … *a*_*j*_}: the *j* bin centers of the histogram of log-10 PaxDb abundances with *a* > *t*
- {*d*,, *d*_1_ *d*_2_, … *d*_*j*_}: the density corresponding to those bin centers

A kernel density estimate *K* was fit to the log10-transformed plasma abundance values using a Gaussian kernel with *σ* = 0.2 and was subtracted from the scaled semi-Gaussian distribution to estimate a function proportional to the density of the probability distribution on abundances for imputed proteins: *h*(*x*) = *αg*(*x*) − *K*(*x*) The function *h*(*x*) was evaluated at 500 abundance values spread equally in base-10 logspace between log10 abundance *log*_10_(*A*_*max*_) − 12 and log10 abundance *log*_10_(*A*_*max*_). Any points where *h*(*x*) evaluated to less than zero were set to zero. A continuous probability distribution was fit to this lattice of sample points using linear interpolation and then normalized such that the total probability of the distribution was 1. The abundances of the 16,017 proteins in the UniProt database not represented in the processed PaxDb dataset were set to random samples from the aforementioned distribution. The resulting abundances were converted to molar fraction estimates by dividing each abundance by the sum of all abundances.

#### Imputation of Protein Abundances (Cell-line)

Abundances were imputed for proteins in the human protein sequence database not represented in the modeled cell-line sample (<u>Protein Abundance Database Processing</u>). This process resulted in a “complete” cell-line sample containing 20,235 proteins with 10 orders of magnitude in dynamic range of abundance. The “complete” cell-line sample was modeled as an adjusted skewed-normal distribution on log10-transformed abundances:

- *g*(*x*) = 2.45 ∗ *skewnorm. pdf*(*x*|*a* = −2.12, *μ* = 4.5, *σ* = 2.55)
- where *skewnorm. pdf* is the probability density function of the skewed normal distribution

A kernel density estimate *K* (Gaussian kernel, *σ* = 0.2) was fit to the log10-transformed abundances of all entries in the processed PaxDb database for the cell-line sample. The function *h*(*x*) was evaluated at 500 abundance values spread equally in base-10 logspace between log10 abundance *log*_10_(*A*_*max*_) − 10 and log10 abundance *log*_10_(*A*_*max*_). Any points where *h*(*x*) evaluated to less than zero were set to zero. A continuous probability distribution was fit to this lattice of sample points using linear interpolation and then normalized such that the total probability of the distribution was 1. The abundances of the 11,923 proteins in the UniProt database not represented in the processed PaxDb dataset were set to random samples from the aforementioned distribution. The resulting abundances were converted to molar fraction estimates by dividing each abundance by the sum of all abundances.

#### Depleted Plasma Sample

To model a plasma sample where the most abundant proteins were depleted from the sample (e.g., using a commercially-available affinity column), the abundances of the top-20 most abundant proteins in the imputed plasma sample (see <u>Imputation of Protein Abundances (Plasma)</u>) were reduced by 99% and the abundances renormalized to sum to 1 to serve as an estimate of molar fraction.

#### Simulating Protein Deposition

Deposition of a sample containing *n* proteins of abundances {*a*_1_ *a*_2_, *a*_3_, … *a*_n_} on a chip was modeled as a multinomial distribution. The protein abundances were normalized to sum to 1 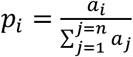. To determine the counts of each protein deposited on a chip with *N* spots, a random sample was made from the multinomial distribution parameterized with probabilities {*p*_1_ *p*_2_, *p*_3_, … *p*_n_} and *N* trials.

#### Simulation of Binding Data

For each sample type (cell, plasma, depleted plasma), binding was simulated for five technical replicate chips. The 300 affinity reagents used for binding targeted the first 300 optimal targets (see <u>Selection and Evaluation of Optimal Affinity Reagent Trimer Targets</u>) and used the binding model described in <u>Biosimilar Affinity Reagent Model</u> with a surface non-specific binding rate of 0.001. To simulate random variation in binding replicate-to-replicate, the binding probabilities of affinity reagents were perturbed for each replicate using the method described in <u>Application of Noise to Affinity Reagent Binding Probabilities</u> with fractional mean absolute deviation 0.1. Binding for each chip was then simulated as described in <u>Simulation of Stochastic Affinity Reagent Binding</u>.

#### Decoding of Binding Data

Protein decoding was performed individually for each replicate as described in <u>Protein Decoding</u>. The human FASTA sequence database (<u>Protein Sequence Databases</u>) was used to define protein candidate sequences. The affinity reagent model used for decoding of all replicates was the original affinity reagent set referenced in <u>Simulation of Binding Data</u> prior to application of random noise. The decode algorithm assumed a surface non-specific binding rate of 0.001.

#### Determining a Probability Threshold for Protein Quantification

At a given identification probability threshold *p*_*t*_, proteins in samples may be quantified by counting the number of identifications for that protein in the decoding output with probability *p* > *p*_*t*_. However, if the probability threshold is set too low, many false positive identifications may occur resulting in low quantitative specificity. If the probability threshold is set too high, false negative identifications may occur, resulting in low quantitative sensitivity. For each replicate chip analyzed, decoding results were processed with probability thresholds: *log*(*p*) = {0, −1 × 10^−20^, −1 × 10^−16^, −1 × 10^−14^, −1 × 10^−12^, −1 × 10^−11^, −1 × 10^−10^, −1 × 10^−9^, −1 × 10^−8^, −1 × 10^−7^, −1 × 10^−6^, −1 × 10^−5^, −1 × 10^−4^, −1 × 10^−3^, −1 × 10^−2^, −0.1, −0.2, −0.3}.

For each threshold evaluated:

- For every unique protein identified at least once in the dataset:
  - Compute the number of reported identifications for the protein that were true positive (i.e., correct identifications) and false positives (i.e., spots incorrectly identified as the protein)
  - Compute the specificity of quantification for this protein: 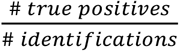
  If the protein has specificity < 0.9, label it as “non-specific identification”
- Compute the “non-specific identification rate”: the fraction of proteins that fall into the “non-specific identification” class

The lowest threshold value resulting in a non-specific identification rate < 0.1% for every replicate analyzed was used for downstream quantification analyses.

#### Quantitative Statistics

After thresholding by identification probability, the following statistics were computed for each analysis:

- Specificity of protein identification was computed as described in <u>Determining a Probability Threshold for Protein Quantification</u>.
- Proteins with at least 1 identification in a given replicate were deemed “identified” in that replicate.
- Proteome coverage for a replicate was the percentage of proteins identified at least once in the replicate among all proteins present in the sample.
- Reproducibility of quantification (CV%) for a protein across replicates was computed using the number of counts for that protein in each replicate: 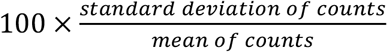. Proteins not identified in a replicate were assigned count 0.

## Acknowledgements

The authors thank Jamie Sherman for helpful discussions on alternative decoding strategies and treatment of non-binding events, Taryn E. Gillies for help with accurately representing epitope locations in the protein workflow figure, Hunter B. Boyce for helpful discussion on strategies for modeling noise in affinity reagent characterization and suggestion of a probit noise model, Patrick McCauley for support with cloud-deployment of decode algorithm, Greg Kapp for helpful suggestions on visually depicting the PrISM workflow, Cristina Zavaleta for ongoing support, Katrina Woolcock from Life Science Editors for major contributions to the writing of the manuscript, Duygu Koldere Vilain from Life Science Editors for major contributions to figures presented in the manuscript, and the broader team at Nautilus Biotechnology for their ongoing critical input to the presented research.

## Author Contributions

PM conceived the presented idea. JE, PM conceived and designed analyses. JE, DD, PM, SP performed analyses. JE, AK, VL, DD, SP, PM developed analysis tools. JE took the lead in writing the manuscript with input from PM. All authors provided critical feedback and helped shape the research, analysis and manuscript.

## Supplementary Figures

**Supplemental Figure 1.**
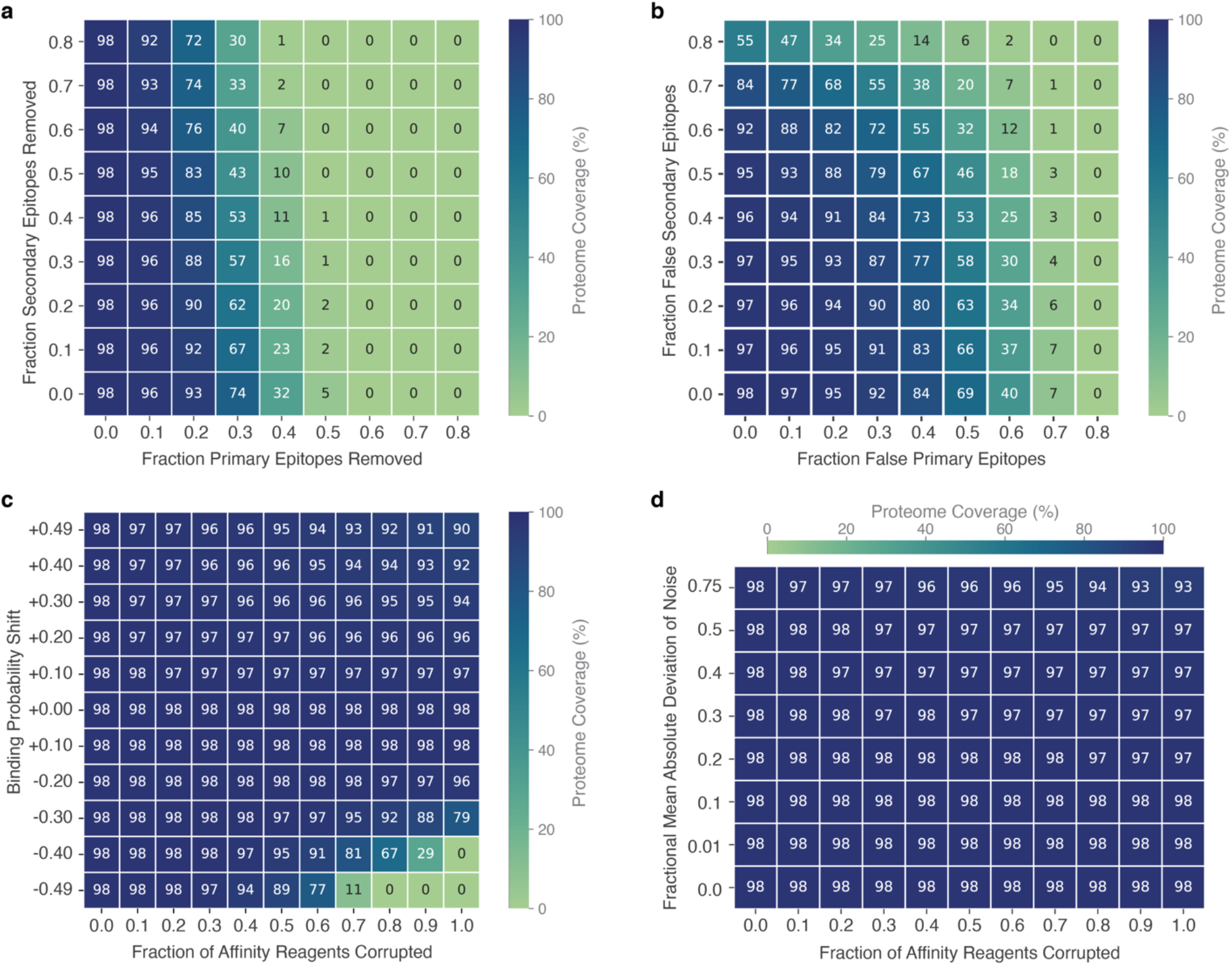
Impact of mischaracterization of affinity reagent binding on proteome coverage for **a)** varying fraction of unknown high-affinity (primary) epitope targets and low-to-medium affinity (secondary) epitope targets, **b)** varying fraction of false high-affinity (primary) and low-to-medium affinity (secondary) epitope targets identified, **c)** systematic measurement error in binding probability with varying fraction of the 300 total affinity reagents impacted by the corruption, and **d)** random measurement error in binding probability with varying fraction of the 300 total affinity reagents impacted by the corruption. All coverage measurements are the average over 5 technical replicates.

**Supplemental Figure 2.**
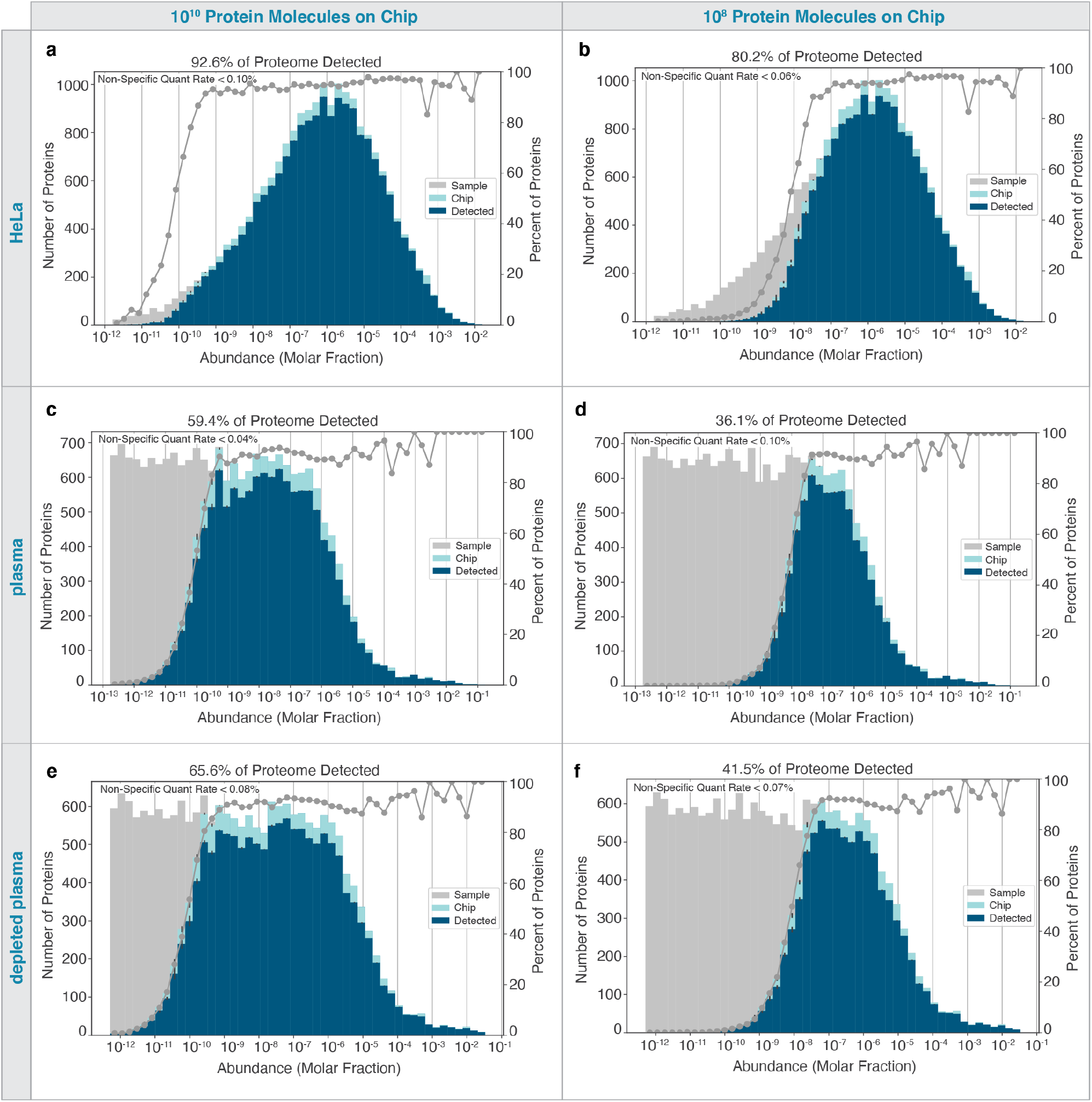
Distribution of protein abundance among proteins in the sample, deposited on the chip, and quantified by PriSM in HeLa cells measured on a 10^10^ landing pad **a)** and 10^8^ landing pad **b)** chip, plasma measured on a 10^10^ landing pad **c)** and 10^8^ landing pad **d)** chip, and a depleted plasma measured on a 10^10^ landing pad **e)** and 10^8^ landing pad **f)** chip. Histogram counts for each group are averaged over 5 simulated replicate experiments. The displayed non-specific quant rate is the maximum percentage of proteins observed in any replicate with poor quantification (>10% signal arising from false identifications). The percent of proteins in the sample quantified is shown as a gray line. Mean proteome coverage is the percent of proteins present in a sample that are detected by PrISM (averaged across the 5 replicates). Error bars indicate standard deviation.

**Supplemental Figure 3.**
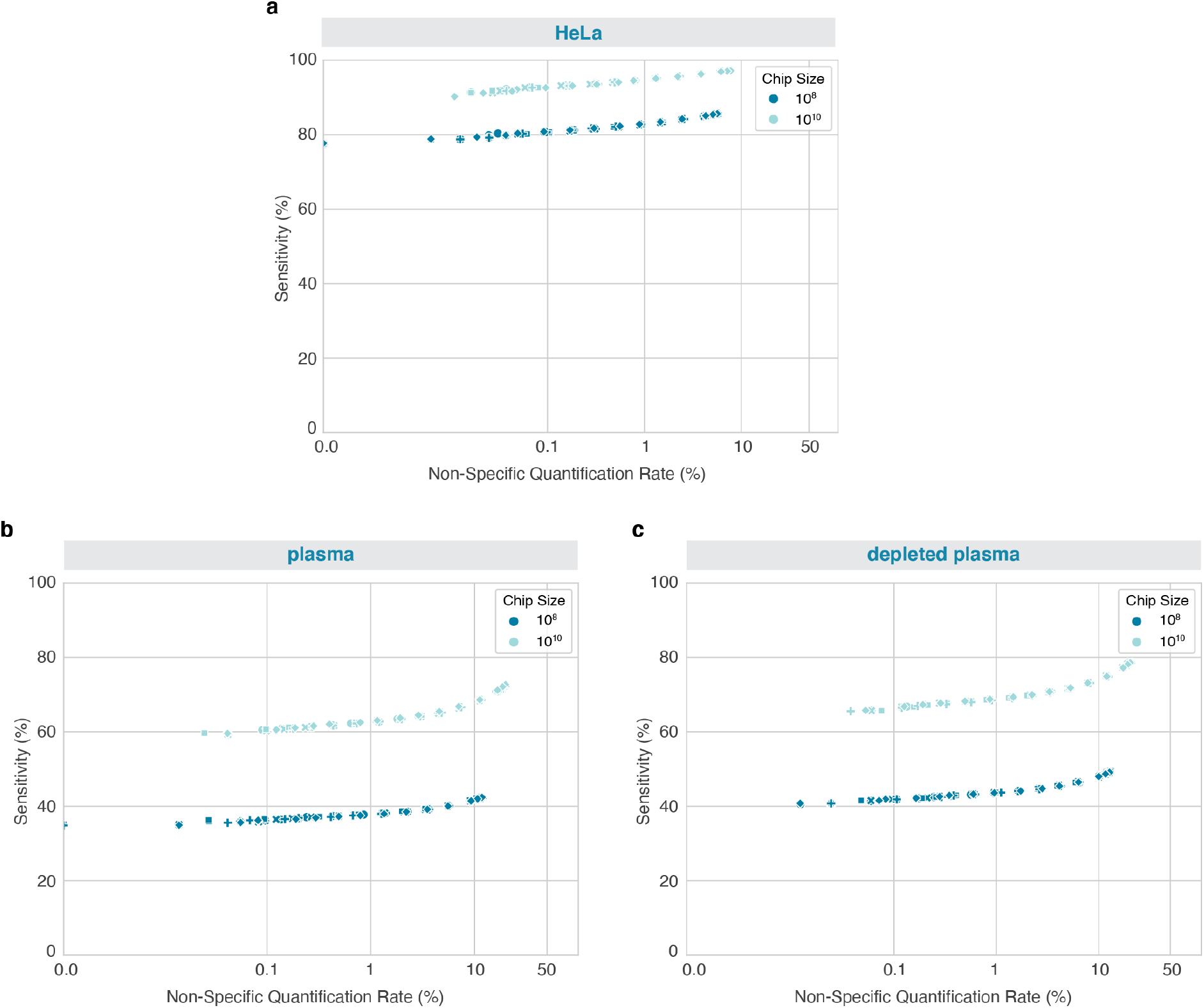
PrISM sensitivity and specificity for HeLa cell line **a)**, plasma **b)**, and depleted plasma **c)**. The probability threshold required for protein identification by PrISM was varied*. A low threshold results in higher sensitivity (proteins quantified) but also a higher rate of non-specific quantification (signals where 10% or more of identifications are false). A point is plotted indicating these metrics for each threshold assessed for each of 5 replicate samples (shown as varying shapes). Simulations were performed with datasets comprising 10^10^ landing pads and 10^8^ landing pads. *18 probability thresholds were assessed: log(threshold)= 0, −1 × 10^−20,^, −1 × 10^−16^, −1 × 10^−14^, −1 × 10^−12^, −1 × 10^−11^, −1 × 10^−10^, −1 × 10^−9^, −1 × 10^−8^, −1 × 10^−7^, −1 × 10^−6^, −1 × 10^−5^, −1 × 10^−4^, −1 × 10^−3^, −1 × 10^−2^, −0.1, −0.2, and −0.3

**Supplemental Figure 4.**
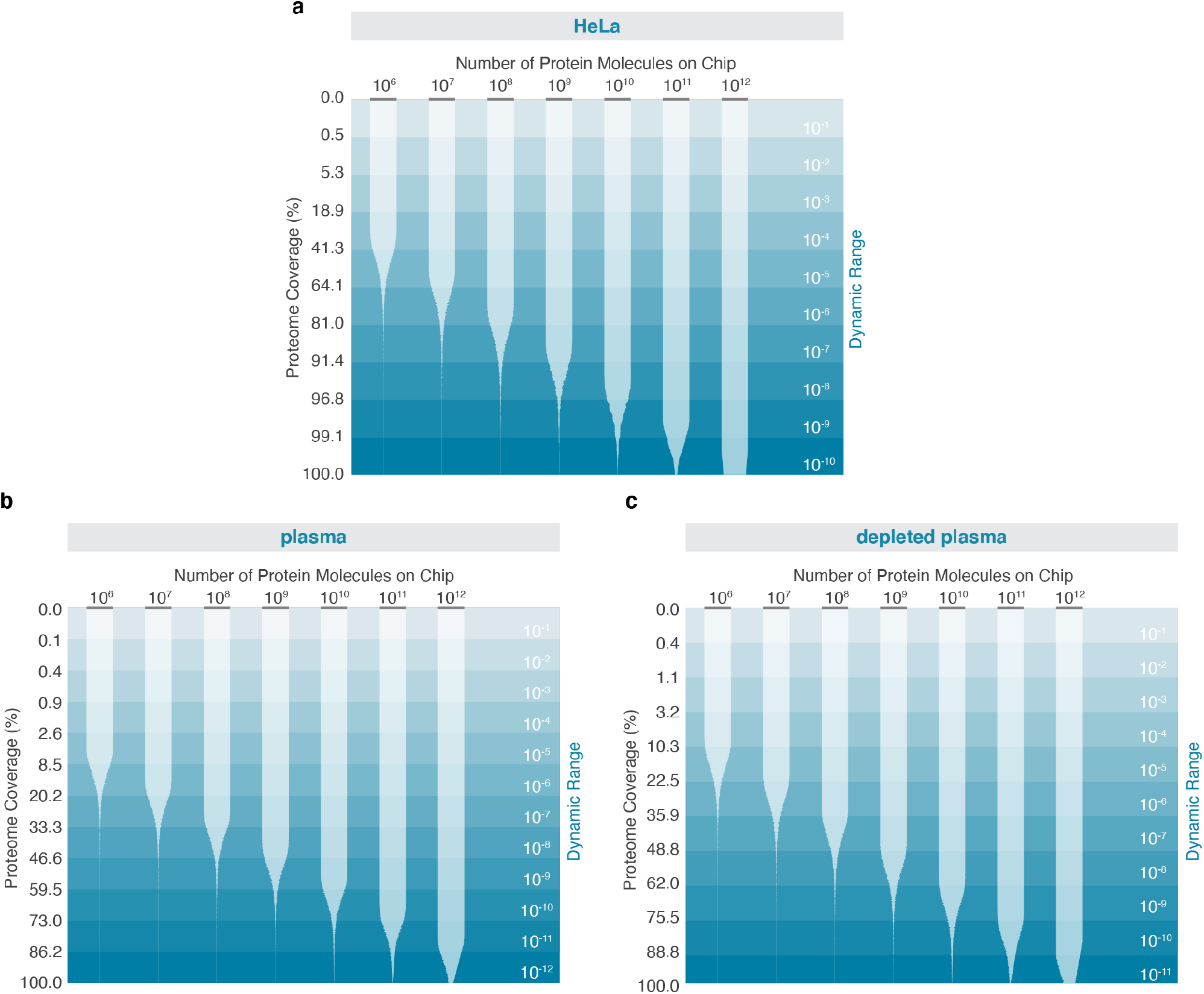
Dynamic range in abundance of proteins deposited on chips of varying size for **a)** HeLa cells, **b)** plasma, and **c)** depleted plasma. Data are plotted in order of decreasing protein abundance from top to bottom. Dynamic range is the protein abundance divided by the most abundant protein in sample. The outer width of the contours indicates the percentage of proteins at that abundance deposited on the chip (one or more copies). The inner width of the contours indicates the percentage of proteins at that abundance detected by PrISM. Percentages are computed over a rolling window of 51 proteins. Horizontal gray bars indicate 100%.

**Supplemental Figure 5.**
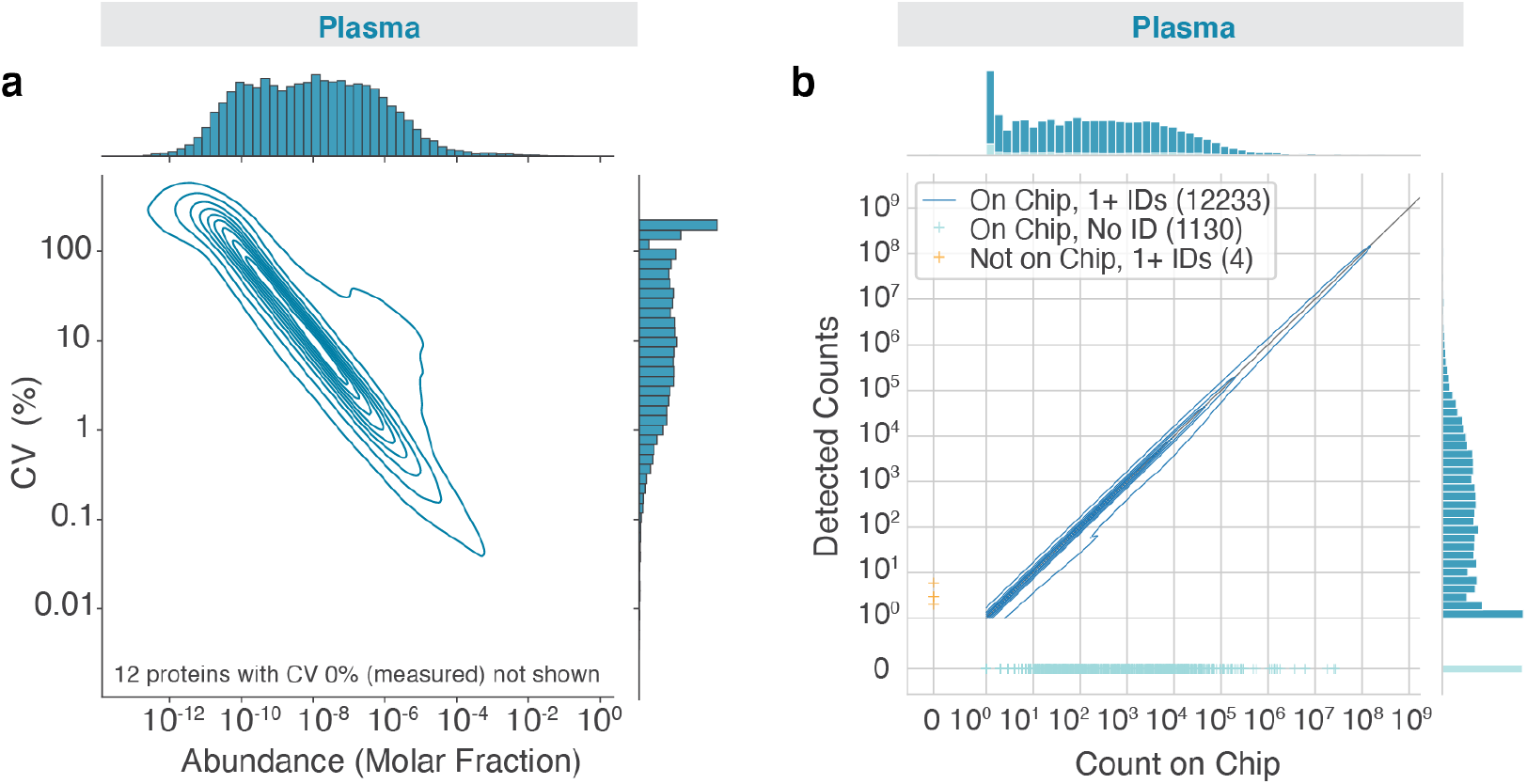
The reproducibility **a)** of quantification (coefficient of variation computed across five replicates) compared to protein abundance for a plasma sample as contour plots (density iso-proportional contours) with marginal histograms and the **b)** concordance of quantity of proteins (number of copies identified) measured by PrISM with true count of protein on chip for a single experimental replicate of plasma as contour plots (density iso-proportional contours) with marginal histograms.

**Supplemental Figure 6.**
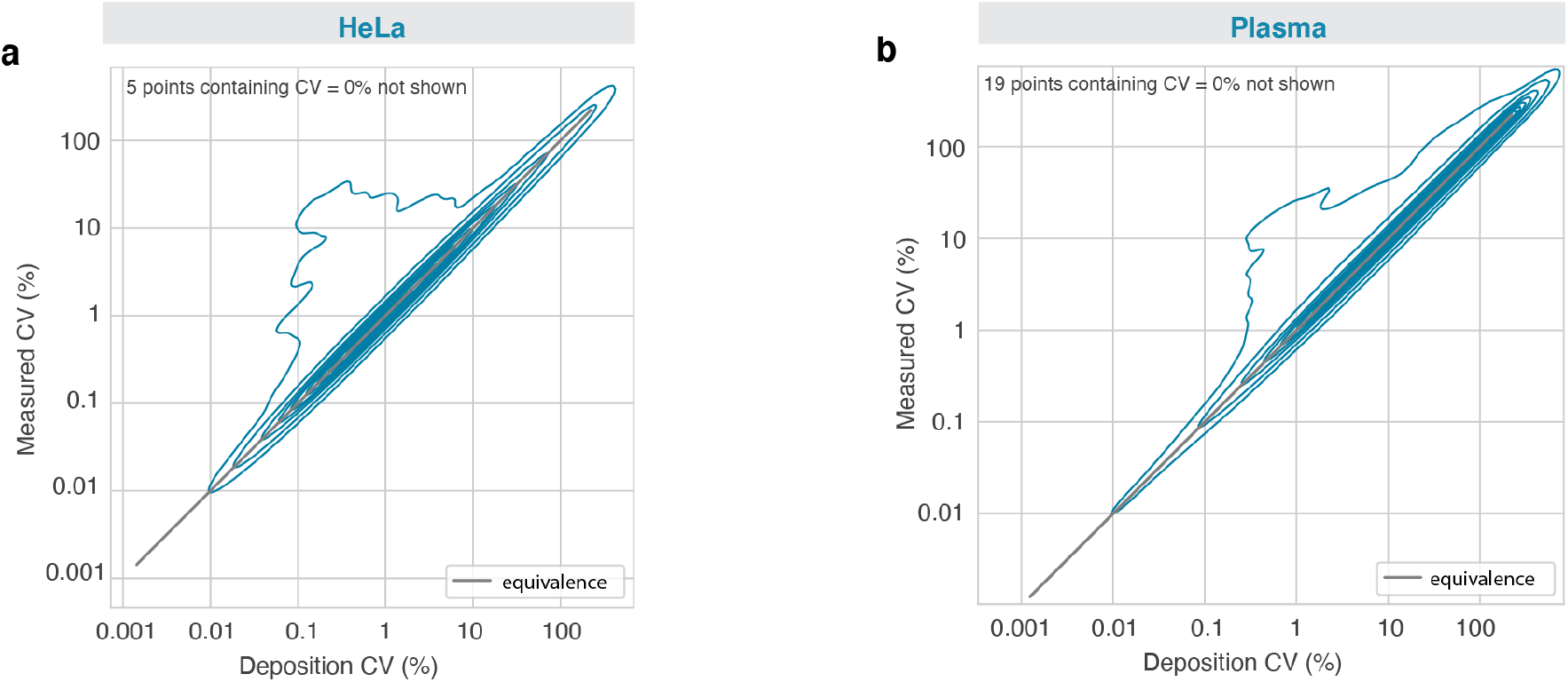
The reproducibility of protein deposition and protein quantification across 5 replicates compared for HeLa cell samples **a)** and plasma **b)** measured on chips with 10^10^ landing pads. Protein quantity deposited is the total count of a protein that was successfully deposited on the chip. Protein quantity measured is the number of times the protein was identified by PrISM. The CV (%) of each of these quantities across the 5 replicates is computed for each unique protein detected in the sample and plotted using a contour plot to demonstrate the concordance of variation in protein counts deposited with variation in protein counts measured.

**Supplemental Figure 7.**
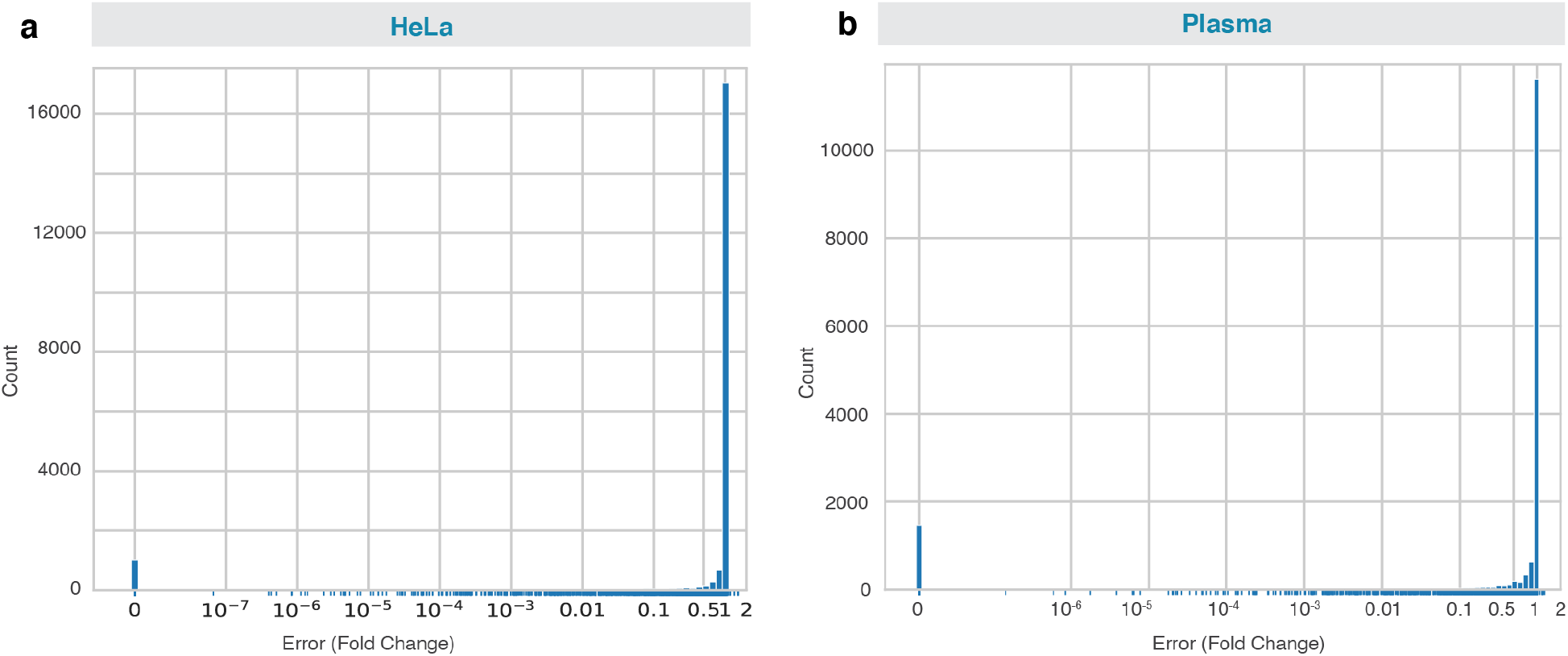
Fold-change measurement error distribution for proteins detected in **a)** HeLa cell line and **b)** plasma samples measured on 10^10^ landing pad chips. Fold change error is the count of protein copies detected by PrISM divided by copies of protein deposited on the chip. Copies detected and copies deposited are averaged across 5 replicates measured.

**Supplementary Table 1.** Affinity reagent characteristics

Affinity reagent characteristics
| Epitope Type | Number of Epitopes per Affinity Reagent | Number of Cycles for 90% Coverage | Number of Affinity Reagents Bound Per Protein (mean) | Number of Affinity Reagents Bound Per Protein (std dev) | % Landing Pads Lit Per Affinity Reagent (mean) | % Landing Pads Lit Per Affinity Reagent (std dev) |
| --- | --- | --- | --- | --- | --- | --- |
| Dimer | 1 | 110 | 40.30728935 | 19.55502431 | 36.64299032 | 14.84719629 |
| Dimer | 2 | >2000 | 1102.618384 | 400.209814 | 55.1309192 | 15.78071197 |
| Trimer | 1 | 410 | 12.53679269 | 11.32908544 | 3.057754314 | 2.335865074 |
| Trimer | 2 | 250 | 14.44343958 | 12.2351679 | 5.777375834 | 3.636218605 |
| Trimer | 5 | 130 | 17.64635532 | 12.8720439 | 13.57411948 | 6.871051851 |
| Trimer | 10 | 100 | 23.70956264 | 14.70880367 | 23.70956264 | 10.00887062 |
| Trimer | 20 | 120 | 44.13743514 | 21.73416995 | 36.78119595 | 14.22655523 |
| Trimer | 25 | 150 | 62.95384235 | 28.56932135 | 41.96922823 | 14.86283342 |
| Trimer | 30 | 220 | 100.8525822 | 42.68880344 | 45.8420828 | 15.54123936 |
| Trimer | 40 | 690 | 361.7437608 | 136.1278615 | 52.426632 | 16.54646207 |
| Tetramer | 1 | >2000 | 3.552409192 | 3.996546384 | 0.17762046 | 0.215077644 |
| Tetramer | 2 | >2000 | 6.561551767 | 6.901374069 | 0.328077588 | 0.344402837 |
| Tetramer | 5 | 1350 | 10.71208302 | 10.60025175 | 0.793487631 | 0.730490262 |
| Tetramer | 10 | 760 | 11.43864591 | 10.92199034 | 1.505084988 | 1.22358617 |
| Tetramer | 20 | 430 | 12.63602669 | 11.43641187 | 2.938610857 | 2.210826628 |
| Tetramer | 25 | 370 | 13.20800593 | 11.69248829 | 3.569731333 | 2.62810432 |
| Tetramer | 30 | 320 | 13.50155671 | 11.71124183 | 4.219236471 | 3.073673392 |
| Tetramer | 40 | 260 | 14.38838646 | 12.00967132 | 5.533994792 | 3.901460484 |

## References

1. Blume, J.E. et al. Rapid, deep and precise profiling of the plasma proteome with multi-nanoparticle protein corona. Nat Commun 11, 3662 (2020).

2. Clark, D.J. et al. Integrated Proteogenomic Characterization of Clear Cell Renal Cell Carcinoma. Cell 180, 207 (2020).

3. Aebersold, R. et al. How many human proteoforms are there? Nat Chem Biol 14, 206–214 (2018).

4. Omenn, G.S. et al. Research on the Human Proteome Reaches a Major Milestone: >90% of Predicted Human Proteins Now Credibly Detected, According to the HUPO Human Proteome Project. J Proteome Res 19, 4735–4746 (2020).

5. Adhikari, S. et al. A high-stringency blueprint of the human proteome. Nat Commun 11, 5301 (2020).

6. Alfaro, J.A. et al. The emerging landscape of single-molecule protein sequencing technologies. Nat Methods 18, 604–617 (2021).

7. Restrepo-Perez, L., Joo, C. & Dekker, C. Paving the way to single-molecule protein sequencing. Nat Nanotechnol 13, 786–796 (2018).

8. Keifer, D.Z. & Jarrold, M.F. Single-molecule mass spectrometry. Mass Spectrom Rev 36, 715–733 (2017).

9. Rissin, D.M. et al. Single-molecule enzyme-linked immunosorbent assay detects serum proteins at subfemtomolar concentrations. Nat Biotechnol 28, 595–599 (2010).

10. Swaminathan, J. et al. Highly parallel single-molecule identification of proteins in zeptomole-scale mixtures. Nat Biotechnol (2018).

11. Swaminathan, J., Boulgakov, A.A. & Marcotte, E.M. A theoretical justification for single molecule peptide sequencing. PLoS Comput Biol 11, e1004080 (2015).

12. Kolmogorov, M., Kennedy, E., Dong, Z., Timp, G. & Pevzner, P.A. Single-molecule protein identification by sub-nanopore sensors. PLoS Comput Biol 13, e1005356 (2017).

13. Chang, L. et al. Single molecule enzyme-linked immunosorbent assays: theoretical considerations. J Immunol Methods 378, 102–115 (2012).

14. Beyer, M. et al. Combinatorial synthesis of peptide arrays onto a microchip. Science 318, 1888 (2007).

15. Anderson, N.L. & Anderson, N.G. The human plasma proteome: history, character, and diagnostic prospects. Mol Cell Proteomics 1, 845–867 (2002).

16. Wang, M. et al. PaxDb, a database of protein abundance averages across all three domains of life. Mol Cell Proteomics 11, 492–500 (2012).

17. Nagaraj, N. et al. Deep proteome and transcriptome mapping of a human cancer cell line. Mol Syst Biol 7, 548 (2011).

18. Eriksson, J. & Fenyo, D. Improving the success rate of proteome analysis by modeling protein-abundance distributions and experimental designs. Nat Biotechnol 25, 651–655 (2007).

